# A severe bottleneck impacted the genomic structure of egg-eating cichlids

**DOI:** 10.1101/2023.05.17.541056

**Authors:** Minami Imamoto, Haruna Nakamura, Mitsuto Aibara, Ryo Hatashima, Takehiko Itoh, Masato Nikaido

## Abstract

The explosive adaptive radiation of Haplochromine cichlids in Lake Victoria, East Africa, produced 500 endemic species within only 15,000 years. A paedophage or an egg-eater is considered a unique example of trophic adaptation. Many field studies reported that more than 200 cichlids have extinct due to the upsurge of Nile perch, a carnivorous species introduced to the lake in the 1950s. Especially, piscivorous cichlids like paedophages were critically damaged by Nile perch predation. Here, we performed a genome-wide evolutionary study of the paedophages in Lake Victoria to understand their past demographic events and phylogenetic relationships. We discovered evidence of a recent, short-period, and severe bottleneck in a paedophage “matumbi hunter”. Interestingly, the signature of a strong bottleneck, as observed in matumbi hunter, was not detected in other species including paedophagus species. In addition, it was revealed that the population size of matumbi hunter started to decline 30 years ago and recover from 20 to 10 years ago, corresponding to the time of both disappearance and resurgence of Lake Victoria Haplochromines were reported. Although population structure analyses showed that matumbi hunter is composed of a unique genetic component, phylogenetic analyses supported its strong monophyly with other paedophagus species. These results suggest that the paedophages originated only once in Lake Victoria followed by the decline of genetic diversity in matumbi hunter. This study succeeded to demonstrate the demographic events triggered by invasive species and associated genomic consequences of the unique trophic group, promoting a holistic understanding of adaptive radiation.

## Introduction

A comprehensive study to evaluate a population bottleneck from polymorphic data is still drawing attention in conservational biology. Theoretical study expected that population bottleneck, resulting in a loss of genetic diversity in a population caused by demographic contraction, generally induces accumulation of deleterious mutation caused by the dominance of genetic drift unlike natural selection, which may result in further population decline or extinction (Ohta 1992). Hence, estimating the time, period, and magnitude of bottleneck in endangered species using genetic data is the one effective solution to consider a proper conservational strategy.

Lake Victoria in East Africa experienced a drastic environmental shift induced by multiple anthropogenic reasons during the twentieth century. One major factor is the rapid upsurge of Nile perch (*Lates niloticus*), a giant carnivore fish, which was introduced to Lake Victoria during the 1950s to 1960s with the purpose to fulfill the increased commercial demand (Fryer 1972; Hauser et al. 1998; Pringle 2005; Mwanja and Mwanja 2008). Nile perch rapidly became the dominant species in the lake, consequently endangering endemic Haplochromine cichlids (Kudhongania and Cordone 1974; Ogutu-Ohwayo 1990; Witte, Goldschmidt, Wanink, et al. 1992; Ogutu-Ohwayo 1993). Haplochromine cichlids in Lake Victoria are known as experienced explosive adaptive radiation, forming 700 endemic species classified into 12 trophic groups (Kaufman 1992; Witte, Goldschmidt, Wanink, et al. 1992; Witte, Goldschmidt, Goudswaard, et al. 1992; Seehausen 1996). On the contrary, more than 200 endemic Haplochromines were estimated as extinct after the expansion of the Nile perch (Ribbink 1987; Ogutu-Ohwayo 1990; Witte, Goldschmidt, Wanink, et al. 1992; Seehausen 1996). Many empirical and observational studies confirmed demographic changes in Haplochromines after the upsurge of Nile perch such as the decline of population size (Kaufman 1992; Witte, Goldschmidt, Goudswaard, et al. 1992; Wanink et al. 2008; Witte et al. 2008; McGee et al. 2015; van Rijssel et al. 2015; Kishe-Machumu et al. 2017; Natugonza et al. 2021)

In 12 trophic groups of Haplochromines, piscivores, a trophic group mainly consuming fish, are considered as experienced a population decline (Witte, Goldschmidt, Goudswaard, et al. 1992; Seehausen 1996; McGee et al. 2015). However, these studies paid major attention to ’fish eaters’ as a representative of piscivores but not ’paedophages (egg-eater)’, a trophic group that feeds eggs and fries by stealing them from female buccal (Greenwood 1959). Here, we specifically focused on *Haplochromis* sp. “matumbi hunter” (matumbi hunter) as an ideal candidate to test population bottleneck. Matumbi hunter is a paedophage distributed in a limited area of Mwanza Gulf, the Southern part of Lake Victoria (Fig. 1a and b). In the comparison of an olfactory receptor gene (*V1R2*) sequences for Lake Victoria Haplochromines, matumbi hunter had a unique allele (V8) out of 20 alleles discovered in the study, implying their poor genetic diversity (Nikaido et al. 2014). Field studies also reported decreased catches of matumbi hunter, coinciding with when Haplochromines started to disappear, inferring that they experienced population contraction (field observation by MA; Seehausen 1996). Thus, we speculated that the matumbi hunter, like the consequence of other piscivores, suffered a population decline due to either the competition with Nile perch or direct predation by Nile perch, resulting in a loss of genetic diversity.

**Figure 1.**
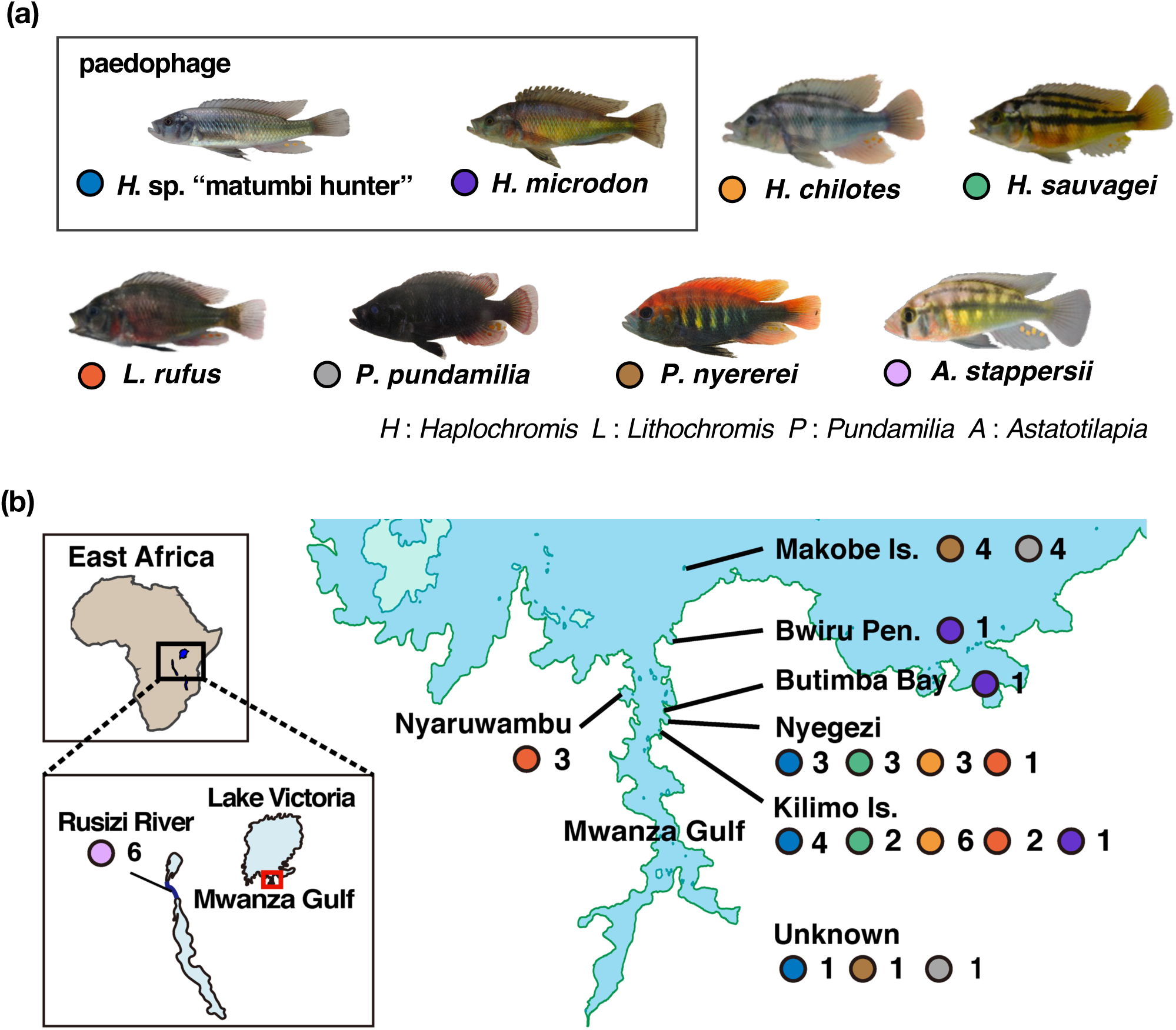
Sampling information and localities of seven Haplochromine cichlids endemic to Lake Victoria and *A. stappersii* the Congolese lineage. (a) Pictures of mainly analyzed eight species. Colored circles next to species names correspond to sampling locations on the map (b). A photo of *A. stappersii* retrieved from Meier et al., 2017. (b) Sampling localities of all individuals in (a). The area marked by a red square in the bottom left map represents the location of Mwanza Gulf Lake Victoria and the map of the enlarged Mwanza gulf is shown on the right. The number of samples per species obtained in each sampling locality is shown next to a circle colored by species, corresponding to labels in (a). Samples without locality information are noted as unknown. Further sampling information and references are written in Table S1.

This study aimed to comprehensively understand the evolutionary history of paedophages in Lake Victoria by mainly focusing on matumbi hunter. We carried out a population structure analysis of 97 species from Lake Victoria and two lineages previously inferred as ancestral lineages of Lake Victoria species flock (Meier, Marques, et al. 2017). While the genetic components were shared among members of paedophages to some extent, matumbi hunter was apparently separated from the other Haplochromines. Next, statistics comparison provided multiple signatures of an intense, short-term, and recent bottleneck in the matumbi hunter, which correspond with when the massive extinction of Haplochromines was reported. We also conducted genome-wide phylogenetic analyses using multiple methods, showing the strong monophyly of paedophages. Consequently, the severe loss of genetic diversity by the upsurge of Nile perch can explain the apparent distinctness of genetic components of matumbi hunter from their closely related paedophagus species. This study newly reports genomic consequences of population decline in paedophagus Haplochromines instigated by the ecological competition with invasive predators.

## Result

### The distinct genetic structure of matumbi hunter in Lake Victoria Haplochromines

To comprehensively understand the genetic structures of Lake Victoria Haplochromines, we carried out genome-wide population structure analyses by ADMIXTURE, PCA, and pairwise *F*_ST_ (Fig. 2).

**Figure 2.**
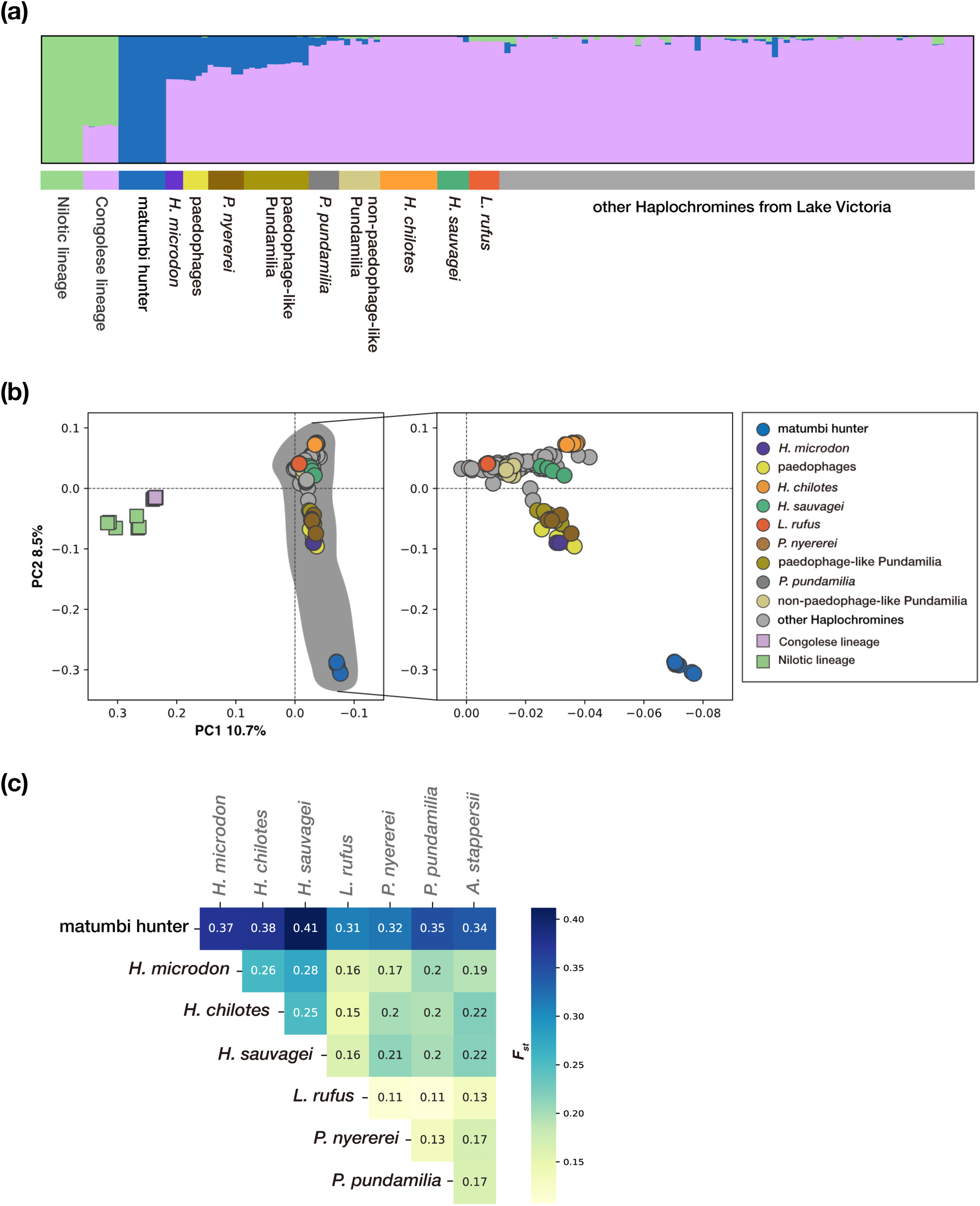
Population structure analyses revealed shared genetic components among paedophages and paedophage-like Pundamilia and elevated differentiation of matumbi hunter. Samples are labeled with grouping name abbreviations: Nilotic lineage (*Astatotilapia bloyeti*, *A. paludinosa*, *Haplochromis gracilior*, and *Thoracochromis pharyngalis*), Congolese lineage (*A. stappersii*), paedophages (*L. parvidens*, *L. melanopterus*, *L. cryptodon*), paedophage-like Pundamilia (*Pundamilia* sp. “big blue red”, *P*. *igneopinis*, *P*. sp. “nyererei-like”, *P*. sp. “orange”, and *P*. sp. “pundamilia-like”), and not-paedophage-like Pundamilia (*P*. sp. “all red”, *P*. *azurea*, *P*. sp. “blue giant”, *P*. sp. “large red deepwater”, *P*. *macrocephala*, and *P*. sp. “pink anal “). (a) ADMIXTURE (K=3) for 97 species from Lake Victoria and their ancestral lineages (Nilotic and Congolese lineages). Labels of all samples are listed in the column “Label in ADMIXTURE” in Table S1. (b) PCA for 97 species from Lake Victoria with two ancestral lineages as an outgroup (left) and without an outgroup (right). Haplochromines endemic to Lake Victoria are shaded as gray in the left figure. The PC1 axis is transposed in descending order for graphical purposes for both figures. The contribution rate for each PC is written on the axis. (c) A heatmap of pairwise weighted *F_ST_* (x-axis versus y-axis) in a total of 28 pairs for 8 populations, a dataset that will be used for statistical comparison in a later section. Boxes with *F*_ST_ values are colored depending on the value of *F_S_*_T._ For example, the *F*_ST_ between matumbi hunter and *H. microdon* is 0.37.

Firstly, we conducted an ADMIXTURE analysis for 97 Haplochromines inhabiting Lake Victoria to compare the genomic components (Fig. 2a, Fig. S1a, Table S1). Species from two lineages: Congolese lineage (*Astatotilapia stappersii*) and Nilotic lineage *(A. bloyeti*, *A. paludinosa*, *H. gracilior*, and *Thoracochromis pharyngalis*) were included in the analysis as an outgroup. The result in K=3 was considered the most reliable K with the lowest CV error rate (Fig. 2a, Fig. S1b). We observed three genetic components: 1) Green observed in both ancestral lineages, 2) Pink observed in Congolese and all Lake Victoria cichlids except matumbi hunter, and 3) Blue observed in paedophages (Fig. 2a). While matumbi hunter was comprised of solely a blue, the large proportion of Lake Victoria cichlids was dominated by a pink, suggesting that matumbi hunter is genetically distinct from other Lake Victoria cichlids. We also carried out PCA with the same dataset used for ADMIXTURE (Fig. 2b, Fig. S2). PC1 represented differences by drainage systems, separating samples by Congolese, Niloticus, and Lake Victoria lineages, whereas PC2 separated matumbi hunter and others, suggesting that matumbi hunter cluster could be isolated from the Lake Victoria cluster including other paedophages. According to the admixture and clustering, we discovered that 13 *Pundamilia* species used in this study were divided into two groups depending on the ratio of paedophage components (pink in Fig. 2a). We call *Pundamilia nyererei* and five *Pundamilia* species (*P*. sp. “big blue red”, *P*. *igneopinis*, *P*. sp. “nyererei-like”, *P*. sp. “orange”, and *P*. sp. “pundamilia-like”) as “paedophage-like Pundamilia”, and *P*. *pundamilia* and six *Pundamilia* species (*P*. sp. “all red”, *P*. *azurea*, *P*. sp. “blue giant”, *P*. sp. “large red deepwater”, *P*. *macrocephala*, and *P*. sp. “pink anal”) as “non-paedophage-like Pundamilia.” Paedophage-like Pundamilia was closely located to the matumbi hunter cluster in PCA (Fig. 2b, Fig. S2). In additional clustering analysis with a dataset including only *Pundamilia* species, paedophage-like and non-paedophage-like Pundamilia formed independent clusters (Fig. S3).

To evaluate the relative magnitude of differentiation between matumbi hunter and other Lake Victoria cichlids, we calculated pairwise *F*_ST_ for all pairs in eight populations (Fig. 2c). Except for matumbi hunter, *F*_ST_ values between Lake Victoria populations ranged from 0.11 to 0.28, and between Lake Victoria and Congolese lineage (*A. stappersii*) ranged from 0.13 to 0.22. Meanwhile, matumbi hunter showed elevated *F*ST values ranging from 0.31 to 0.41, even *F*ST=0.37 with *H. microdon*, the other paedophagus species.

Three population structure analyses were consistent in that the matumbi hunter population is genetically differentiated even from a sympatrically distributing paedophage, while some genetic components were shared among paedophages.

### Signatures of a recent, short-term, strong bottleneck in matumbi hunter population

To further evaluate the distinct genetic structure of matumbi hunter, we observed indicative statistics of genetic diversity and demography. Firstly, we calculated nucleotide diversity (*π*, the mean pairwise differences per base pair) in a 10 KB non-overlapping window (Fig. 3a, Table S2). Matumbi hunter showed the lowest average *ρε* in eight populations, suggesting its poor genetic diversity compared to other species. We calculated inbreeding coefficient *F* by individual sample and then grouped and averaged them according to species (Fig. 3b, Table S2). The elevated *F* in matumbi hunter population was yielded (*F*_mean_=0.49), implying that mating between closely related individuals was frequently occurring, resulting in an excess number of homozygous sites. *H. microdon*, *H. chilotes* and *H. sauvagei* also showed relatively high *F* (0.23, 0.22, 0.25, respectively), while *L. rufus, P. nyererei, and P. pundamilia* exhibited *F* less than 0.1. We observed a negative value of *F* for *A. stappersii*. We also estimated the decay of pairwise linkage disequilibrium (LD) by comparing the squared genotypic correlation coefficient *r*^2^ of pairwise SNPs, located at a distance less than 10 KB (Fig. 3c, Table S2). The increased *r*^2^ in distantly located SNPs in matumbi hunter means that recombination and novel mutation have been less accumulated because of the recent bottleneck. Meanwhile, *L. rufus* and *A. stappersii* showed relatively low *r*^2^ in all ranges of SNP distances.

**Figure 3.**
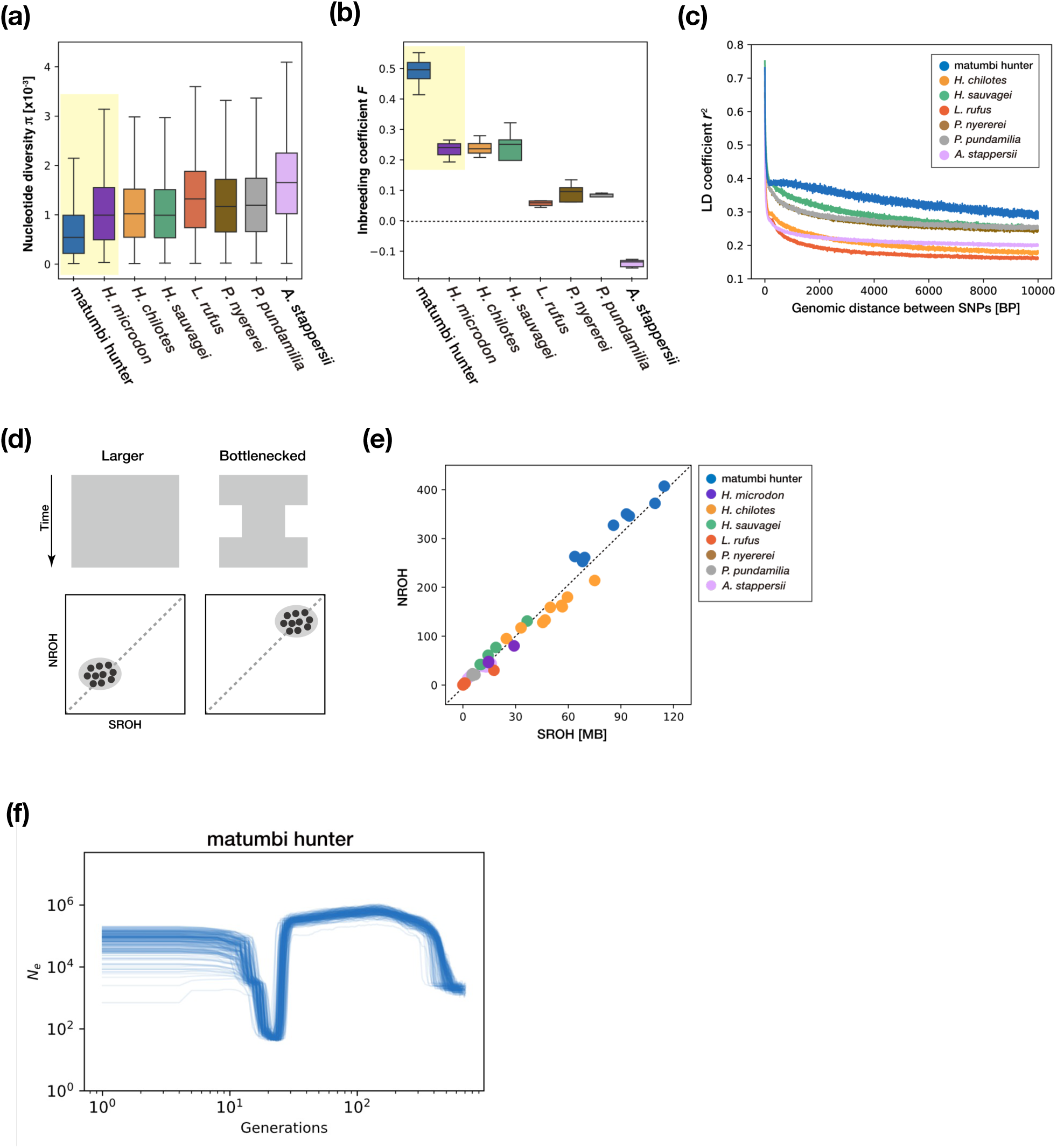
Signatures of the recent, short-term, and strong bottleneck event in the matumbi hunter population. (a) 10 KB windowed nucleotide diversity (*p*) across a genome calculated by species. (b) Averaged inbreeding coefficient (*F*) by population. We calculated *F* for each sample, then averaged them by population. (c) Comparison of linkage disequilibrium (LD) decay plotted LD coefficient (*r*^2^) of SNPs having a distance within 10 KB. *H. microdon* has been excluded from this analysis because an insufficient number of samples per species may cause a bias in estimating *r*^2^. (d) Expected correlation between the sum total length of ROH (SROH) and the total number of ROH (NROH) under certain demographic history (larger (left) and bottlenecked populations (right)). Theoretically, a short and small number of ROH regions (both SROH and NROH are smaller) will be detected in a large and expanded population, whereas long and large numbers of ROH (both SROH and NROH are larger) can be observed in a small and bottlenecked populations (Ceballos et al. 2018) (e) Observed correlation of SROH and NROH. Each plot corresponds to one sample. Homozygous regions lasting more than 150 [KB] were defined as ROH regions. (f) Changes in effective population size (*N*_e_) in matumbi hunter in the past 700 generations. The GONE estimate was repeated 200 times, and all results were plotted.

Next, we obtained genome-wide runs of homozygosity (ROH, contiguous homozygous blocks in the genome), inference to grasp holistic demographic events by observing a correlation between the sum total length of ROH (SROH) and the total number of ROH (NROH). All matumbi hunter samples showed an extended SROH and excessive NROH corresponding to example ROH patterns, strongly suggesting the past bottleneck event (Fig. 3d and e). We further calculated Tajima’s *D* which is used to examine the occurrence of demographic events such as population bottleneck (Tajima 1989). Although we did not observe a significant elevation of Tajima’s *D* in matumbi hunter, a variance of Tajima’s *D* was higher than in other populations (Table S2, Fig. S4).

To directly compare demography among populations, we assessed trajectories of effective population size (*N*_e_) in the last 700 generations by the GONE program (Fig. 3f, Fig. S5). We confirmed the population decline in matumbi hunter started around 30 generations ago, then recovered from 20 to 10 generations ago (Fig. 3f). Defining the generation time of cichlid as 1 year and the sampling year of matumbi hunter as 2018, the bottleneck in matumbi hunter started in 1988, and recovery began around 1998, coinciding with the time when Seehausen (1996) reported them in the publication for the first time. In addition, population decline during the post-Nile perch era has been also observed for *H. microdon*, *H. chilotes*, and *H. sauvagei* (Fig. S5a, b, d, respectively), and they showed a tendency of weak bottleneck in previous statistics comparison (Fig. 3a-c).

### Phylogenetic trees inferred by five methods agreed on the monophyly of paedophages

To elucidate the phylogenetic relationship between matumbi hunter and other haplochromines, we carried out a phylogenetic analysis in five methods considering different theories to estimate a topology. Previous studies struggled to infer phylogenetic relationships of Lake Victoria haplochromines because of their low genetic differentiation due to the recent radiation, only providing star-like topologies so far (Samonte et al. 2007; Wagner et al. 2013; McGee et al. 2020). In this study, phylogenetic trees were constructed in five methods: three coalescence-based methods ((a) ASTRAL-III, (b) SVDquartet, and (c, d) SNAPP in Fig. 4) and two maximum-likelihood-based (ML-based) methods ((e) RAxML-NG and (f) IQ-TREE2 in Fig. 4). As a result, the strong monophyly of five paedophagus cichlids (matumbi hunter (Hmat), *H. microdon* (Hmid), *L. parvidens* (Lpar), *L. melanopterus* (Lmel), and *L. cryptodon* (Lcry)) was reconstructed in all estimated trees except for SNAPP. In the SNAPP tree, matumbi hunter was located at the basal position in the Lake Victoria clade (Fig. 4c). However, DensiTree and consensus tree were partially mismatched (Fig. 4c and d). In additional SNAPP trees from un-concatenated datasets, seven out of ten topologies included matumbi hunter in the paedophage clade (Fig. S6). Thus, in addition to the results of population structure inferences (Fig. 2), phylogenetic analyses strongly supported the monophyly of paedophages, implying that paedophages originated only once in Lake Victoria. Apart from the robustly supported paedophage clade, the other topologies were fragile, in that *H. sauvagei* (Hsau), *H. chilotes* (Hchi), and *L. rufus* (Lruf) was monophyletic in coalescence-based trees (Fig. 4a-d), while they were paraphyletic ML-based trees (Fig. 4e and f). Especially, we observed that five samples of *P. nyererei* formed a paraphyly with poor bootstrap values in a RAxML-NG tree (Fig. 4e) despite that they are grouped in the population structure analyses (Fig. S1a; Fig. S3). It is noteworthy that *P. nyererei* was estimated as sister taxa to the paedophage clade in all inferred phylogenetic trees.

**Figure 4.**
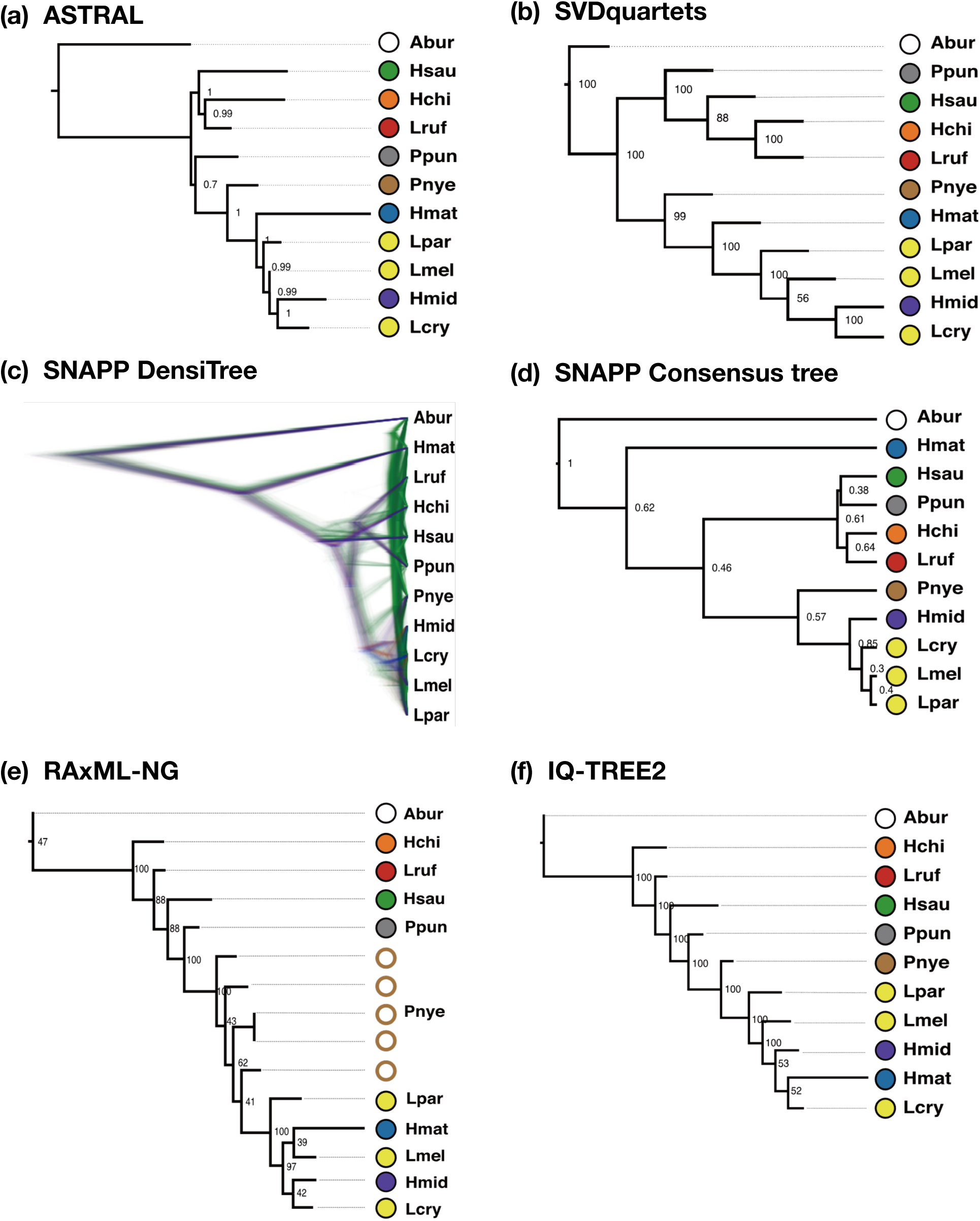
Strong monophyly of paedophage was confirmed by phylogeny estimated by ASTRAL (a), SVDquartets (b), and maximum likelihood methods (RAxML-NG (e) and IQ-TREE2 (f)), except for SNAPP (c, d). *Astatotilapia burtoni* (Abur), a riverine lineage, was rooted as an outgroup for all analyses. Samples are labeled with species name abbreviations: Hmat (matumbi hunter), Hmid (*H. microdon*), Lpar (*L. parvidens*), Lmel (*L. melanopterus*), Lcry (*L. cryptodon*), Pnye (*P. nyererei*), Ppun (*P. pundamilia*), Hchi (*H. chilotes*), Hsau (*H. sauvagei*), Lruf (*L. rufus*). Bootstrap values and posterior probabilities for each node are shown in (b), (e), (f), and in (a), (d), respectively. In (b), (e), and (f), samples formed monophyletic clades by species were collapsed for graphical purposes, except for *P. nyererei* (Pnye) in (d), which showed a paraphyletic topology (brown outlined white circle). (a) A consensus phylogenetic tree obtained by ASTRAL-III, estimated from 7,980 gene trees in a 20 KB sliding window with stepwise 5 KB. (b) Coalescent tree built in SVDquartets with 100 bootstrap replicates. Note that SVDquartets is capable to estimate only the topology but not the other parameters such as branch length. (c) DensiTree was made from concatenated multi-trees generated by 10 independent SNAPP runs, and its consensus tree (d) was estimated by TreeAnnotator in the BEAST 2.0 package. In DensiTree, the more commonly observed trees are colored in the order of blue, red, and green, while the rest are dark green, respectively. (e) Maximum likelihood tree reconstructed by RAxML-NG in 200 bootstraps under GTR+G4 model. (f) The maximum likelihood tree was estimated by IQ-TREE2 with 1000 ultrafast bootstrap replicates under the TVMe+R5 model.

To compare the credibility of inferred phylogenetic relationships in Fig. 4, we carried out a topology weighting by the TWISST program for four arbitrary groups in three ranges of SNP windows (Fig. 5a). For average weightings in all window sizes, topo 1 (out, ((C-I, Ppun), (Pnye, C-Egg))); was the prevalent topology out of 15 possible topologies, which corresponded with the one recovered by SVDquartets (Fig. 5 b, Fig. 4b). Curiously, the major three topologies supported the close relatedness of *P. nyererei* and C-Egg as always recovered in Fig. 4 (Fig. 5b topo1, topo2, and topo3).

**Figure 5.**
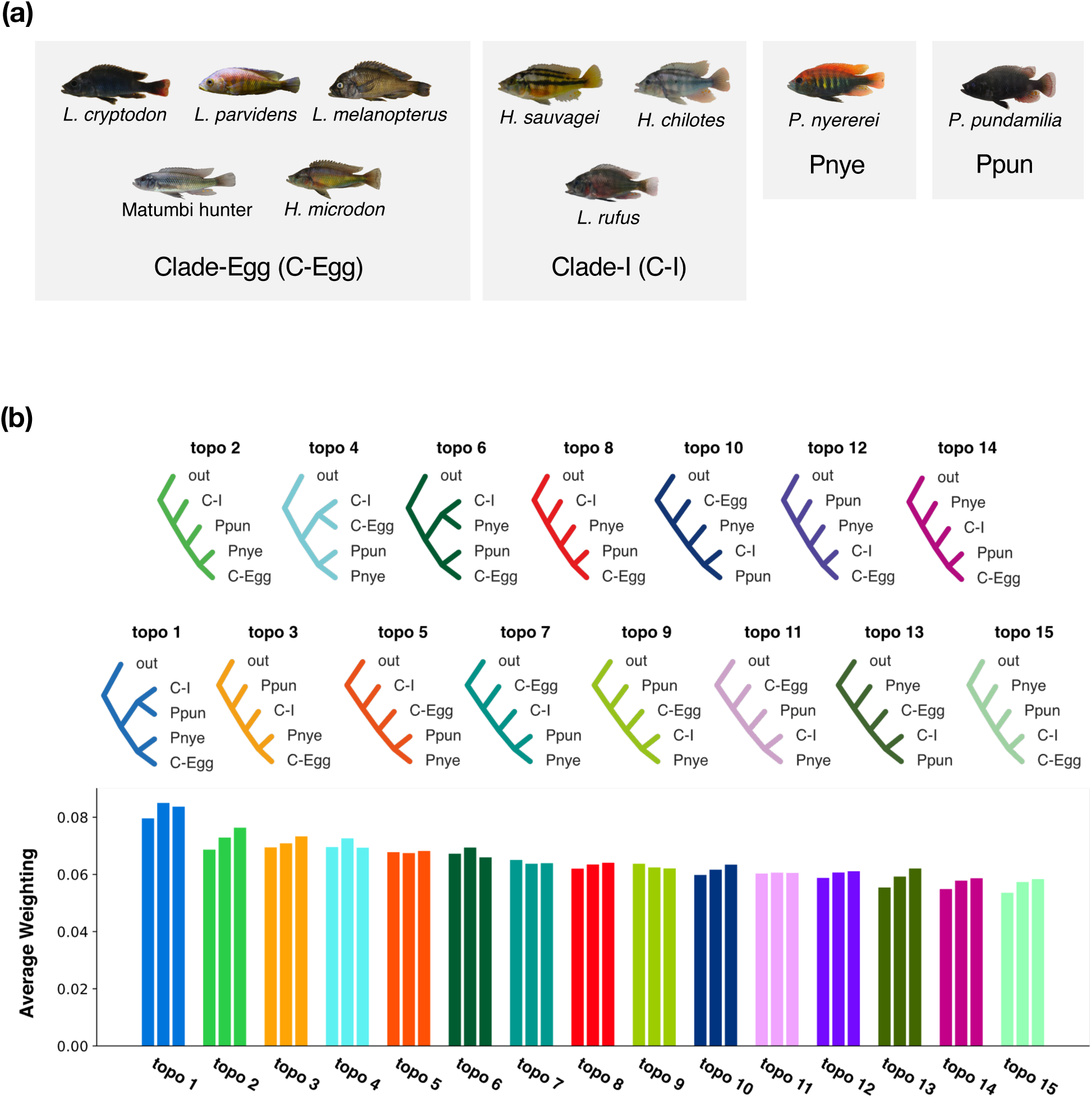
Topology weighting across the genome recovered topologies inferred in phylogenetic analyses as the major topology. (a) Provided groupings for TWISST analysis. According to topologies constructed by phylogenetic analyses (Fig. 4), we defined four topological groups: five paedophages including matumbi hunter as C-Egg, three species often formed monophyletic clade as Clade-I, *P. nyererei*, and *P. pundamilia*. (b) 15 possible topologies for four groups defined in (a) and their average weightings in three window sizes. *Astatotilapia burtoni* was set as Outgroup (out). Three bars for each topology represented by a unique color, are average weightings observed in 50SNP(left), 100SNP(center), and 300SNP(right) windows. Bars are sorted in descending order of average weightings among three windows.

We also inferred the split times for 21 combinations in 7 populations by focusing on that of matumbi hunter (Fig. 6). The average split time between *A. stappersii* and matumbi hunter was about 40,000 years, falling in the range of the average split time between *A. stappersii* and other Haplochromines of Lake Victoria. In addition, the average split times among Haplochromines in Lake Victoria were all about 19,000 years, which approximately corresponds to the radiation of the Lake Victoria Haplochromines (Johnson et al. 2000; Meier, Marques, et al. 2017; Nakamura et al. 2021).

**Figure 6.**
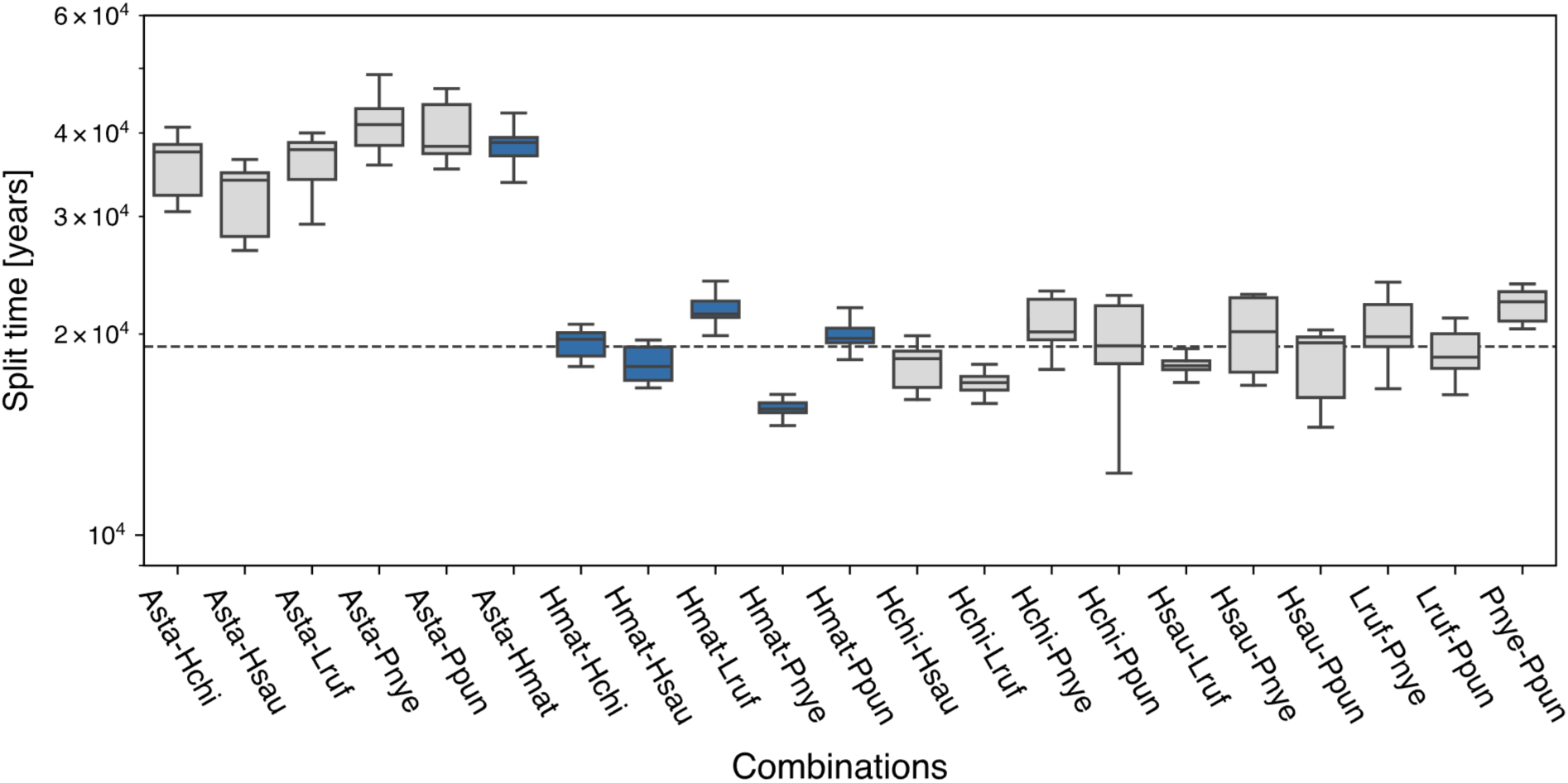
Split times in pairs of studied seven populations proved the co-diversification of Lake Victoria Haplochromines. Samples are labeled with species name abbreviations: Asta (*A. stappersii*), Hmat (matumbi hunter), Hchi (*H. chilotes*), Hsau (*H. sauvagei*), Lruf (*L. rufus*), Pnye (*P. nyererei*), Ppun (*P. pundamilia*). The estimation of split time was repeated 100 times per pair. The per-generation mutation rate was set as 3.5 x 10^-9^ referred by Malilnsky et al., 2018. The mean split time among pairs in Lake Victoria cichlids (19,158 years) was drawn as a dashed line. Split times in pairs of matumbi hunter are colored blue.

## Discussion

### The severe bottleneck in matumbi hunter occurred during the upsurge of Nile perch

The impacts on endemic Haplochromine complex caused by the upsurge of Nile perch have been investigated in aspects of ecological observation and morphological alteration. However, despite their negative effects threatening species diversity, a comprehensive study based on evolutionary genomics evaluating and discussing a demographic event has yet remained to be reported. This study succeeded to demonstrate the magnitude of the population bottleneck and associated genomic consequences in a piscivorous Haplochromine in Lake Victoria.

Evolutionary genomics approaches discovered signatures of poor genetic diversity and population decline in matumbi hunter, concluding as it was a strong, short-term, and recent bottleneck greatly corresponded with the time of massive extinctions of cichlids in the 1980s, in which expansion of Nile perch is considered as the significant factor (Ribbink 1987; Ogutu-Ohwayo 1990; Witte, Goldschmidt, Wanink, et al. 1992; Seehausen 1996). Poor nucleotide diversity in matumbi hunter can be assumed as the consequence of extreme population shrinkage (Fig. 3a, Table S2). In population structure analyses, matumbi hunter was genetically differentiated from other sympatric species, as *F*_ST_ between matumbi hunter and other species was higher than *F*_ST_ in pairs of *A. stappersii*, the early diverged riverine lineage (Fig. 2c). However, at the same time, the phylogenetic analysis didn’t detect any early divergence of matumbi hunter among studied species and matumbi hunter was ingroup of Lake Victoria clade (Fig. 4, Fig. 6). Hence, we assume that the genetic differentiation of matumbi hunter was triggered by a strong bottleneck. The demographic estimation revealed that *N*_e_ in matumbi hunter had started to drastically decrease to less than *N*_e_ =100 in 30 -20 years ago (1988-1998; Fig. 3f), which is the period when the extinction of endemic Haplochromines was becoming a concerning issue (Ogutu-Ohwayo 1990; Kaufman 1992; Witte, Goldschmidt, Goudswaard, et al. 1992). Then 20-15 years ago (1998-2003) was the *N*_e_ recovery phase, and finally, *N*_e_ completely increased to the current size, again coinciding with when the partial resurgence of Haplochromine was reported (Witte et al. 2007; Natugonza et al. 2021). A *N*_e_ in the population contraction phase of bottleneck might reflect an excess number of homozygous loci as seen in an elevated inbreeding coefficient and ROH pattern (Fig. 3d and e, Table S2). Strong LD at distantly located SNPs indicates less frequent recombination on chromosomes, implying a very recent event of a bottleneck (Fig. 3c). Such observed parameters, a recent, short-term, and strong bottleneck may have complicated inferring demographic insight from global Tajima’s *D* across the genome but instead, local Tajima’s *D* by locus was highly variable in matumbi hunter (Fig. S4, Table S2). Actually, the simulation study expected that elevated variance of Tajima’s *D* across the genome indicates the fluctuation of the population size (Nakamura et al. 2018).

Interestingly, although we succeeded to detect a bottleneck event in matumbi hunter, a signature of such bottleneck effect was not observed in *H. microdon*. Matumbi hunter and *H. microdon*, another paedophage analyzed in this study, are having mutual ecomorphological features; a relatively slender body rarely observed in rock-dwelling cichlids, and a sympatric distribution inhabiting Mwanza Gulf, unlike other paedophages mainly inhabiting outside of the gulf. Thus, we expected that if the upsurge of Nile perch was the crucial factor of a bottleneck in matumbi hunter, *H. microdon* should also have experienced a population contraction. *H. microdon* also presented low nucleotide diversity and high homozygosity among analyzed populations, but their bottleneck was not as harsh as the one in matumbi hunter (Fig. 3a and b). In a demographic inference, a short-term population decline seemed to happen in *H. microdon*, though, we were unable to determine the consensus ages of any demographic event from fluctuating estimation by each replicate (Fig. S5a). In fact, it is important to note that we failed to include *H. microdon* in some analyses, such as LD coefficient, demography, and split time due to a lack of sampling numbers. Some phenotypic characteristics are only seen in matumbi hunter such as a gray coloration which is hardly observed among either paedophages or rock-dwelling cichlids, and inclined outer teeth in lower jaws unlike other paedophages (Seehausen 1996). Such morphological differences might have contributed a severe bottleneck specifically in matumbi hunter.

### The phylogenetic origin of trophic adaptation in paedophage

This study is the first example performed large-scale evolutionary genomics of paedophages in Lake Victoria, leaving signatures explaining parallel and independent trophic adaptation within Lake Victoria regions. It is common that similar eco-morphologies are mutually observed throughout the great three lakes in East Africa, which are considered as parallelly and independently evolved at each lake (e.g. stripe patterns on a body (Urban et al. 2022) and hypertrophied lip (Kocher et al. 1993; Darrin Hulsey et al. 2018)). While no paedophagus species have been reported in Lake Tanganyika, several species in Lake Malawi (McKaye and Kocher 1983; Dierickx and Snoeks 2020), 24 species including undescribed species in Lake Victoria (Witte et al. 1976; Greenwood 1980; Witte and van Oijen 1990; Seehausen 1996), and 6 species in the surrounding satellite lakes (1 species in Lake Kivu (Snoeks, 1994) and 5 species in Lake Edward (Snoeks 1994; Vranken et al. 2019)) have been observed. Especially, Vranken et al. (2019) reported the anatomical similarities between *H. microdon* and *H. occultidens* a paedophage inhabiting Lake Kivu, providing an intuitive example of the parallel evolution of paedophage in the Lake Victoria region.

Our phylogenetic analyses including matumbi hunter and four representative paedophages using multiple methods proved that they mostly form monophyletic “the paedophage clade” in all reconstructed topologies (Fig. 4). In addition, ADMIXTURE analysis suggested the existence of paedophage-specific genomic regions, which may have been sustained among paedophages (Fig. 2a, Fig. S1a). Hence, we would like to propose that the distinct population structure of paedophages explains the single origin of paedophages in the Lake Victoria region. We also confirmed an elevated ratio of shared genetic components among paedophages despite the massive loss of genetic diversity in matumbi hunter, implying genomic regions relating to their unique eco-morphologies are highly conserved. The phylogenetic relationship and population structure of paedophages proposed in this study hopefully may promote a holistic understanding of the mechanism triggered by explosive trophic radiation and associated ecological specialization that occurred in Lake Victoria, and the genetic mechanism controlling feeding behavior.

This study also ensured the close relatedness between paedophage and paedophage-like Pundamilia, implying that their relationship is the key to understanding ecological diversification. In ADMIXTURE, paedophage-like Pundamilia contained genomic components spreading among paedophages (Fig. 2a), and *P. nyererei* was frequently recovered at the root of the paedophage clade in phylogenetic analyses (Fig. 4, Fig. 5). McGee et al. (2020) built a big-picture phylogeny by mtDNA and partial genomic regions and IBD network using Lake Victoria cichlids, inferring not only the close relatedness of paedophages but also their close phylogenetic relationship with some *Pundamilia* species, which we called as paedophage-like Pundamilia in the current study. Hence, we would like to conclude that paedophage-like Pundamilia is the closest lineage of paedophages, and ecological adaptation might have happened in the common ancestor of these lineages. Interestingly, Seehausen (1996) reported “*Pundamilia* sp. ’nyererei paedophage’” from field observation at the western Mwanza Gulf, the newly discovered and undescribed species. If we consider the close phylogenetic relationship between paedophage and *P. nyererei*, individuals with such hybrid morphs perhaps were sustaining features from both lineages, reflecting a state in the middle of divergence. Interestingly, the genome-wide relatedness with paedophages mentioned above was observed in a limited *Pundamilia* species which we called paedophage-like Pundamilia. As we visualized a distinct population structure of paedophage-like versus non-paedophage-like Pundamilia in PCA, there might be two groups within the genus *Pundamilia* (Fig. S3)

### Understanding the genetic background underlying diverse ecologies

A large-scale population structure analysis using about a hundred cichlids from Lake Victoria succeeded to capture ecology-distinct patterns of admixture represented as strong allele share among paedophages (Fig. 2a, Fig. S1a). A genetic admixture after drastic fluctuation of the lake water level is considered the major factor that fueled explosive adaptive radiation in Lake Victoria (Salzburger 2018). Takeda et al. (2013) performed population structure and haplotype network analyses in over ten populations of Lake Victoria cichlids using mtDNA and microsatellites, despite low differentiation levels between populations, revealing distinct genetic backgrounds by population reflecting eco-morphological variations and demographic history. In Nakamura et al. (2021), a comparative genomics study of the three species of haplochromines revealed population-specific demography, assuming that they had adapted to their own ecologies. Our findings in the current study also succeeded to recover unique genetic background reflecting ecological variation, greatly agreeing with previous studies. Neutrality statistics inferring genetic diversity and demographic events and estimation of fluctuations of *N*_e_ proved the population-specific genomic consequences (Fig. 3, Table S2, Fig. S4-6).

The phylogenetic analysis provided fine-scaled phylogenetic trees with leaving the consensus tree, proving the effectiveness of estimation in multiple methods with genome-wide SNPs of multiple samples per species. Although many studies have been combatted to infer the phylogenetic relationship of Haplochromines in Lake Victoria, a low genetic differentiation among species due to incomplete lineage sorting (ILS) disrupted to infer a consensus phylogenetic tree reflecting true evolutionary history. Theoretically, the most frequently observed topology across the genome in topology weighting should correspond to the species tree reflecting a true evolutionary history, even if there is a partial gene flow between analyzed populations (Martin and Van Belleghem 2017). However, in our topology weighting, the difference in average weightings between topo1 and topo2 was not drastically elevated (Fig. 5b). Topologies except for the topo1 have also been equally observed across the genome, suggesting that the topology is highly differed by the genomic regions thus ensuring the frequent ILS in Lake Victoria Haplochromines. Scherz et al. (2022) succeeded to recover a genus-level phylogeny among Mbuna cichlids in Lake Malawi which has been radiated within less than two million years, using genome-wide SNPs by searching the consensus topology from trees constructed in multiple methods. Thus, this study also constructed phylogenetic trees using five major methods to compare inferred topologies, to avoid a gene tree conflict caused by ILS. Topologies inferred by ASTRAL-III and SVDquartets both coalescent-based methods almost agreed except for the allocation of *P. pundamilia* (Fig. 4a and b). Indeed, demographic simulation dating speciation and gene flow events between *P. pundamilia* and *P. nyererei* from Makobe populations inferred that gene flow occurred at least twice within the last 6,000 years (Meier, Sousa, et al. 2017), thus we should consider the possibility that their admixture events might have complicated phylogenetic inferences, explaining a fluctuated topologies of *P. pundamilia*. Despite high heterozygosity across the cichlid genome, previous phylogenetic studies normally used a single individual per species without performing haplotype phasing. To avoid an issue discarding informative heterozygous sites, we implemented ASTRAL-III, a method known as an effective phylogenetic analysis tool for recently diverged lineages since it uses phased data as an input and considers the assignment of heterozygous sites for the phylogenetic estimation (Zhang et al. 2018), a theoretically suitable approach for Lake Victoria cichlids.

## Conclusion

The current study implemented population genetics and phylogenetics studies for representative Haplochromines in Lake Victoria using large-scale genomic data. We newly proved that matumbi hunter, a paedophage endemic to Lake Victoria, experienced a strong population decline coinciding with the period of the upsurge of Nile perch, and they are currently in the recovery/expansion phase. Evolutionary genomics of five paedophages revealed that they mostly form a strict monophyletic clade, suggesting lake-independent trophic adaptation. We also showed that Pundamilia species can be classified into two groups: the group that shares genetic components with paedophages (paedophage-like Pundamilia), and the one that doesn’t (non-paedophage-like Pundamilia). Paedophage-like Pundamilia diverged at the root of the paedophage clade in phylogenetic analyses, implying ecological adaptation to paedophage has occurred at that divergence. In addition, our strategy to estimate a phylogenetic relationship of recently diverged lineages, which uses multiple samples per species, multiple estimation methods, and phased-genome data allowed us to infer consensus topology in fine resolution. This study would facilitate an understanding of mechanisms that regulate unique feeding behavior and massive trophic adaptation of Haplochromines in Lake Victoria.

## Materials and Methods

### Samples and Resequencing

This study used 158 samples of whole genome sequences in total, including 137 samples sequenced in previous studies obtained from the NCBI database and 21 samples newly re-sequenced in this study (Table S1). Wild males of *Haplochromis* sp. ’matumbi hunter’, *H. microdon*, *H*. *sauvagei*, *H*. *chilotes*, and *Lithochromis rufus* were collected in Mwanza Gulf and its surroundings in Lake Victoria by M.A.. Genomic DNA was extracted from either fin clips or muscles following protocols of DNeasy Blood & Tissue Kit (QIAGEN). After we prepared Paired-end libraries following protocols of TruSeq DNA PCR-Free LT sample Prep Kit, whole genome resequencing was performed by Illumina HiSeq 2500.

### Mapping and Variant Calling

Firstly, we generated a master VCF file containing all samples by the following procedure. The quality of short reads was checked by FastQC (Andrews 2010), and adapter sequences and bad quality reads were removed by fastp (Chen et al. 2018). Raw reads were aligned to the *de novo* assembled genome of *H. chilotes* (Nakamura et al. 2021) by bwa-mem (v. 0.7.17-r1188) (Li 2013). Unique reads with acceptable mapping quality was kept out using the option “f-2 -F2052 -q 30” in samtools (v.1.8) (Li et al. 2009). After extractions of initial variants by bcftools (v. 1.8) (Li 2011), only SNPs with minimum depth=10, maximum depth=60, and minimum mapping quality 60 have remained using “vcffilter” command in vcflib (Garrison et al. 2022). Single VCF files of all individuals were merged by bcftools. We removed all missing sites, insertions, and deletions, sites that deviate from HWE (p-value < 0.001), and non-biallelic sites (using “--max-missing 1”, “--remove-indels”, “--hwe 0.001”, and “--max-alleles 2 --min-alleles 2” commands in vcftools (v. 0.1.16) (Danecek et al. 2011), respectively), remained 622,956,232 neutral SNPs defined as “master VCF file.”

### Datasets Preparation

We prepared the following three VCF datasets from the master VCF file,

1. Population-dataset for population structure estimation
2. Statistics-dataset for genetic statistics calculation
3. Phylogeny-dataset for phylogenetic analysis

keeping only candidate samples and filtering SNPs in different options. Samples included in each dataset are written in the “ds1”, “ds2”, and “ds3” columns in Table S1.

### 1. Population-dataset

Intending to infer allele-share and ancestry among lineages, 97 species endemic to Lake Victoria and species classified as two ancestral groups Congolese and Nilotic lineages were kept as Population-dataset. We kept 155 samples and excluded SNPs with frequency less than 0.01 then converted to PLINK accessible format (“--keep”, “--maf 0.01” and “--plink” commands in vcftools, respectively). SNPs in linkage disequilibrium (LD), defined as SNPs with LD coefficient *r*^2^ >0.1, were estimated and pruned by PLINK v1.9 (Chang et al. 2015) “--indep-pairwise 50 5 0.1” and “--extract” commands, respectively, remaining 1,282,071 SNPs.

### 2. Statistics-dataset

Seven species from Lake Victoria and one ancestral lineage *Astatotilapia stappersii* from Rusizi River, containing more than three samples per population at least were selected to calculate genetic statistics inferring genetic diversity and demography (Fig. 1). 47 samples were kept from the master VCF file using the command “--keep” in vcftools and remained 37,335,868 SNPs.

### 3. Phylogeny-dataset

For revealing the phylogenetic relationship within paedophages and their closely related lineages, five paedophages, five sympatric species, and one early diverged lineage *A. burtoni* were included in Phylogeny-dataset as an outgroup. We kept 45 samples and excluded SNPs with frequency less than 0.03 then converted to Plink accessible format (“--keep”, “--maf 0.03” and “--plink” commands in vcftools, respectively). SNPs in linkage disequilibrium (LD), defined as SNPs with LD coefficient *r*^2^ >0.1, were estimated and pruned by plink “--indep-pairwise 100 5 0.1” and “--extract” commands, respectively, remained 298,674 SNPs.

### Population Structure and Differentiation Study

Population structure and differentiation analysis were performed by following procedures using 1. Population-dataset. We analyzed individual ancestry with ADMIXTURE (v1.3.0) (Alexander et al. 2009), estimating the proportion of components in K=2 to K=8. The K with the lowest cross-validation (CV) error value was selected as the optimal grouping. Eigenvalues and vectors were generated by the PLINK function “--pca” and then principal components with high eigenvalues were plotted to visualize population structure. Pair-wise genetic divergence (Weir and Cockerham weighted *F*_ST_ (Weir and Cockerham 1984)) between eight populations was estimated in a 10,000 bp window by vcftools from 2. Statistics-dataset.

### Population Genomics

Statistics inferring genetic diversity and demographic events were calculated by using 2. Statistics-dataset. Nucleotide diversity (*ρε*) and Tajima’s *D* within a population in a 10,000 bp window and inbreeding coefficient (*F*) within an individual were computed with vcftools functions “--window-pi 10000”, “--TajimaD”, and “--het”, respectively. LD coefficients (*r*^2^) between SNPs located within 10,000 bp were observed by PopLDdecay (Zhang et al. 2019). We additionally prepared a filtered VCF dataset for runs of homozygosity (ROH) analysis, removed SNPs with frequency less than 0.03, and converted to PLINK format (“--maf 0.03” and “--plink” vcftools commands, respectively), then SNPs with LD coefficient *r*^2^ >0.1 have been excluded by “--indep-pairwise 50 5 0.1” PLINK command. The sum and number of ROH within an individual, defined as a homozygous region lasting more than 150 kbp, were inferred by PLINK commands “--homozyg --homozyg-density 5 --homozyg-kb 150 --homozyg-snp 2”.

### Effective population size

We implemented GONE (v1.0) (Santiago et al. 2020) to estimate *N*_e_ in recent generations such as -700 generations, aimed to detect demographic changes in the pre-and post-Nile perch era. To fulfill computational requirements in GONE, we thinned 2. Statistics-dataset by keeping scaffolds longer than 150 KB. The thinned dataset was kept by population and converted to PED and MAP by PLINK, manually replacing scaffold names with ascending integers. To avoid a biased estimation in admixed population, we assigned the maximum value of recombination which showed monodisperse demography (0.05 for matumbi hunter, *H. microdon*, and *H. sauvagei*, and 0.01 for *H. chilotes*, *P. pundamilia*, *P. nyererei*, *L. rufus*, and *A. stappersii*). Effective population size in the last 700 generations has been estimated with an option randomly extracting 22,000 SNPs per scaffold (modified as “NGEN=700”, “NBIN=700”, “maxNCHROM=-99” and “maxNSNP=22000” in INPUT_PARAMETERS_FILE), replicating 200 times to infer the consensus demography.

### Phylogenetic Relationships

#### RAxML-NG, IQ-TREE2, and SVDquartets - 3

Phylogeny-dataset has been converted to phylip-format by vcf2phylip (Ortiz 2019). GTR+G4 was selected as an appropriate model by ModelTest-NG (v. x.y.z) (F louri et al. 2015; Darriba et al. 2020). The maximum likelihood tree was estimated by RAxML-NG (v. 0.9. 0) (Kozlov et al. 2019) with 200 bootstrap replicates. IQ-TREE2 (v. 2.0.3) (Nguyen et al. 2015) has been implemented with 1000 ultrafast bootstrap replicates (Hoang et al. 2018) and TVMe+R5 model estimated b y ModelFinder (Kalyaanamoorthy et al. 2017). To infer the phylogenetic tree on SVDquartets, we converted 3. Phylogeny-dataset to nexus-format by convert_vcf_to_nexus.rb (available at https://github.com/mmats chiner/tutorials/blob/master/species_tree_inference_with_snp_data/src). We run the SVDquartets (Chifman and Kubatko 2014; Chifman and Kubatko 2015) implemented method in PAUP (v 4.0) (Swofford 2003) t o infer the multispecies coalescent tree with 100 bootstrap replicates.

#### ASTRAL-III

As ASTRAL-III (v5.7.8) (Zhang et al. 2018) expects diploid datasets, we additionally prepared a phased-VCF dataset. Before phasing scaffolds, less than 50 SNPs were removed from 3. Phylogeny-dataset to avoid phasing errors. We implemented haplotype phasing by Beagle (v5.2) (Browning et al. 2021) for 50 iterations, and confirmed all heterozygous sites were phased. ASTRAL-III input multi-tree file was created by running raxml_sliding_window.py (available at https://github.com/simonhmartin/genomics_general), containing 7,980 gene trees in 20 KB sliding window with stepwise 5 KB. Finally, we ran ASTRAL-III under default settings, assigning all samples into species, and obtained a consensus phylogenetic tree.

#### SNAPP

To infer species trees under the multispecies coalescent model, the SNAPP program has been implemented using the BEAST 2.0 platform (Bryant et al. 2012; Bouckaert et al. 2019). As SNAPP requires significant computing power to run large datasets, we replicated the analysis with thinned datasets containing randomly selected 1000 SNPs from 3. Phylogeny-dataset. Thinned datasets were converted to nexus format by vcf2phylip. XML files for the BEAST 2.0 run were manually generated in BEAUTi (GUI in the BEAST 2.0 platform). We implemented SNAPP analysis in BEAST 2.0 for 2 million generations, recording topologies every 1000 steps. We confirmed that the effective sample size of each run exceeded 200 by Tracer (Rambaut et al. 2018). Outputs were visualized in three different ways: independently annotated trees (un-concatenated, Fig. S6), DensiTree of concatenated multi-trees of a dataset (Fig.4c), and its consensus tree (Fig.4d). Firstly, we directly inferred consensus trees for each dataset by TreeAnnotator in the BEAST 2.0 platform with 10% burn-in and visualized them in FigTree (v1.4.4) (Rambaut 2006). Multiple trees from datasets were concatenated by LogCombiner in the BEAST 2.0 platform. To grasp the distribution of all inferred trees, multi-trees in concatenated data were drawn in layered by DensiTree (Bouckaert and Heled 2014), colored based on the more commonly observed trees in the order of blue, red, and green, while the rest are dark green, respectively. From multi-trees in concatenated data, the maximum clade credibility tree was determined by TreeAnnotator with 10% burn-in and drawn by FigTree.

### Topology weighting across the genome using TWISST

Topology weightings in multiple window sizes were estimated by TWISST (Martin and Van Belleghem 2017). From the phased-VCF file generated in ASTRAL-III dataset-preparation, we ran by raxml_sliding_window.py to generate three multiple-tree files containing 731216, 366759, and 125123 gene trees in 50-, 100-, and 150 bp-windows, respectively. We defined *Astatotilapia burtoni* as an outgroup and classified other ten species into four groups referring to already estimated phylogenies: five paedophagus species including matumbi hunter as C-Egg, three species frequently formed monophyly as Clade-I, *P. nyererei* as Pnye, and *P. pundamilia* whose separation timing was different by methods as Ppun (Fig. 5a). Topology weightings for 15 possible topologies were estimated for all input gene trees. Finally, consensus topology was determined by comparing average weightings per topology.

### Speciation time

Speciation times for all possible 21 pairs in seven populations were calculated by SMC++ (v1.15.4) (Terhorst et al. 2017). *H. microdon* was excluded from this analysis because we were concerned that the lack of sample size may disturb to an inference of true split time. A VCF dataset prepared in the previous GONE implementation, filtered out scaffolds shorter than 150 KB, was converted into “.smc” format with chunking VCF by populations and scaffolds (“smcpp vcf2smc” command in SMC++). We generated 100 inputs per scaffold for replicated runs by bootstrap_smcpp.py (available at https://github.com/popgenmethods/smcpp/issues/37), implementing *N*_e_ estimations for 100 replicated runs (with the command “smcpp estimate”) by setting the per-generation mutation rate as 3.5 x 10^-9^ (Malinsky et al. 2018). Ranges of generations to calculate *N*_e_ were heuristically determined by software.

Finally, we run to estimate speciation time with the function “smcpp split” in SMC++, setting the per-generation mutation rate as 3.5 x 10^-9^ (Malinsky et al. 2018).

## Acknowledgments

This study was supported by JSPS KAKENHI (20J13861 to H. N., 20KK0167 to M. N.) and JST SPRING (JPMJSP2106 to M. I.). Phylogenetic analyses were partially performed using the NIG supercomputer at ROIS National Institute of Genetics. Whole genome sequence reads of newly reported samples have been deposited to DDBJ Sequence Read Archive (SRA) under the accession ID XXX.

## Author Contributions

M. I. performed population genetic and genomic analyses and wrote the paper. R. H. carried out genomic DNA extraction and purification for whole genome resequencing. T. I. conducted a whole genome resequencing of some samples newly reported in the study. H. N. carried out a bioinformatic analysis processing short-read data to extracted SNPs. H. N., M. A., and M. N. provided their expertise and commented on the paper while M. A. was responsible for sample collection. H. N. and M. N. supervised the study.

## Legends to Figures

## Legends to Tables

**Table S1. Summary information of all genome samples analyzed in the current study.** The “Color” column is shaded by a color representing each population. If the exact locality in Lake Victoria where the sample has been caught is known, noted in the “Locality” column, otherwise written as “unknown”. For samples newly sequenced in this study and previously sequenced in our group, sampled years are written in the “Sampled year” column. If the sample was included in 1. Population-dataset, 2. Statistics-dataset, or 3. Phylogeny-dataset, marked as “+” (“-” if not included) in the “ds1”, “ds2”, and “ds3” columns, respectively. Bin numbers in ADMIXTURE analysis are assigned in the “Label in ADMIXTURE” column. McGee et al. (2020) and Seehausen (1996) defined *Lipochromis cryptodon* and *L*. sp. “velvet black cryptodon” as different species though, out analysis summarized them as one population because no elevated genetic differentiation has been detected between them.

**Table S2. Summary of population statistics to infer signature of a bottleneck.** Mean values of nucleotide diversity, inbreeding coefficient, linkage disequilibrium (LD) coefficient, and Tajima’s *D* including variance, averaged by species, are listed in the table, respectively. LD coefficient in *Haplochromis microdon* was not determined as lack of sampling number will disrupt to infer true statistics.

**Table S3. Summary of mean split times in 21pairs for 7 populations and average split time among analyzed Haplochromines in Lake Victoria.** Samples are labeled with species name abbreviations: Asta (*A. stappersii*), Hmat (matumbi hunter), Hchi (*H. chilotes*), Hsau (*H. sauvagei*), Lruf (*L. rufus*), Pnye (*P. nyererei*), Ppun (*P. pundamilia*). The estimation of split time was repeated 100 times per pair. The per-generation mutation rate was set as 3.5 x 10-9 referred by Malilnsky et al., (2018).

**Figure S1.**
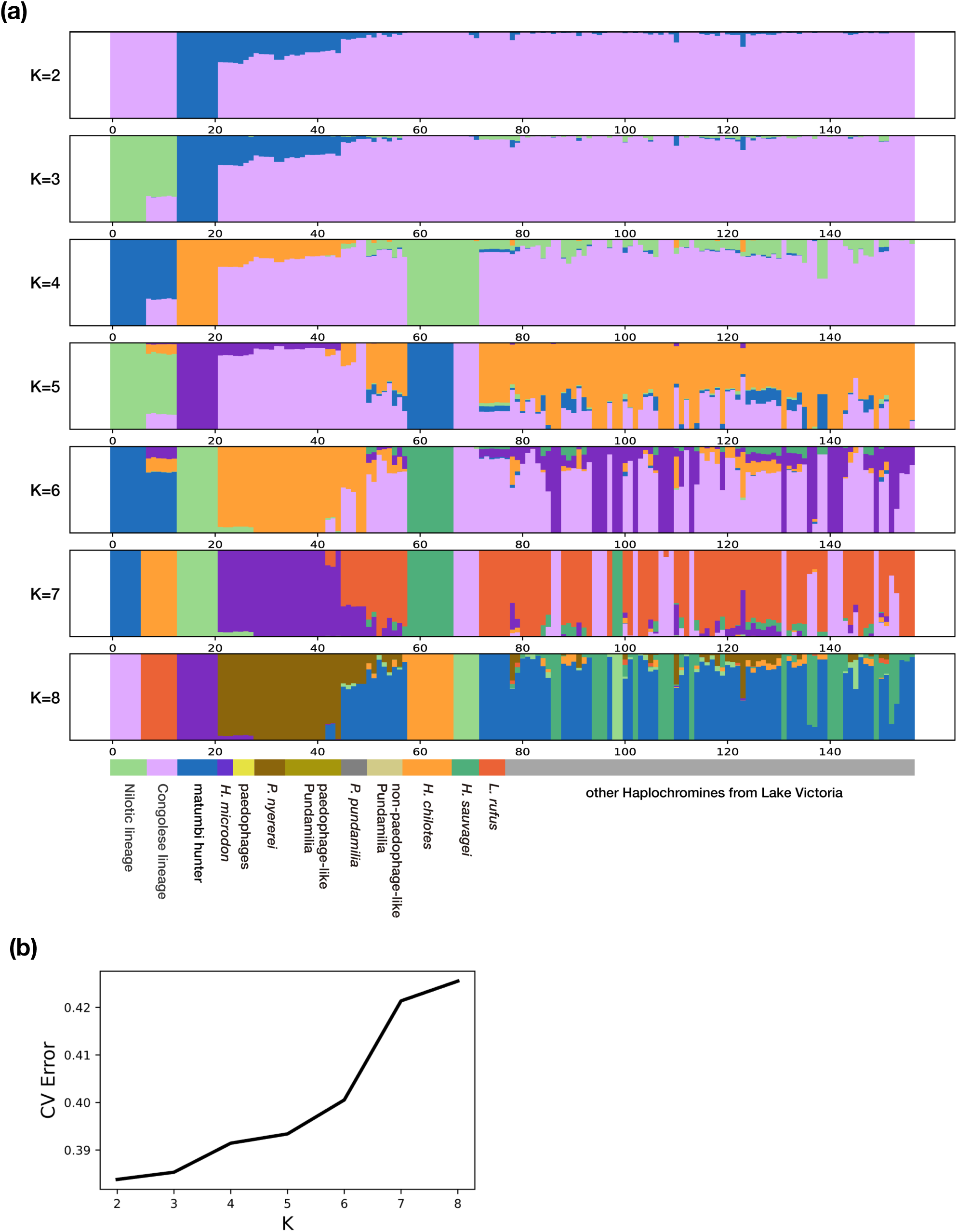
ADMIXTURE in the range of K=2 to K=8 (a) and its cross-validation (CV) errors by K (b). Samples are labeled with grouping name abbreviations: Nilotic lineage (*Astatotilapia bloyeti*, *A. paludinosa*, *Haplochromis gracilior*, and *Thoracochromis pharyngalis*), Congolese lineage (*A. stappersii*), paedophages (*L. parvidens*, *L. melanopterus*, *L. cryptodon*), paedophage-like Pundamilia (*Pundamilia* sp. “big blue red”, *P*. *igneopinis*, *P*. sp. “nyererei-like”, *P*. sp. “orange”, and *P*. sp. “pundamilia-like”), and not-paedophage-like Pundamilia (*P*. sp. “all red”, *P*. *azurea*, *P*. sp. “blue giant”, *P*. sp. “large red deepwater”, *P*. *macrocephala*, and *P*. sp. “pink anal “). (a) Result of ADMIXTURE analyses in K=2 to K=8 for 97 species from Lake Victoria and their ancestral lineages (Nilotic and Congolese lineages). Labels of all samples are listed in the column “Label in ADMIXTURE” in Table S1. (b) CV error comparison. The lower CV error at a K implies a more reliable result.

**Figure S2.**
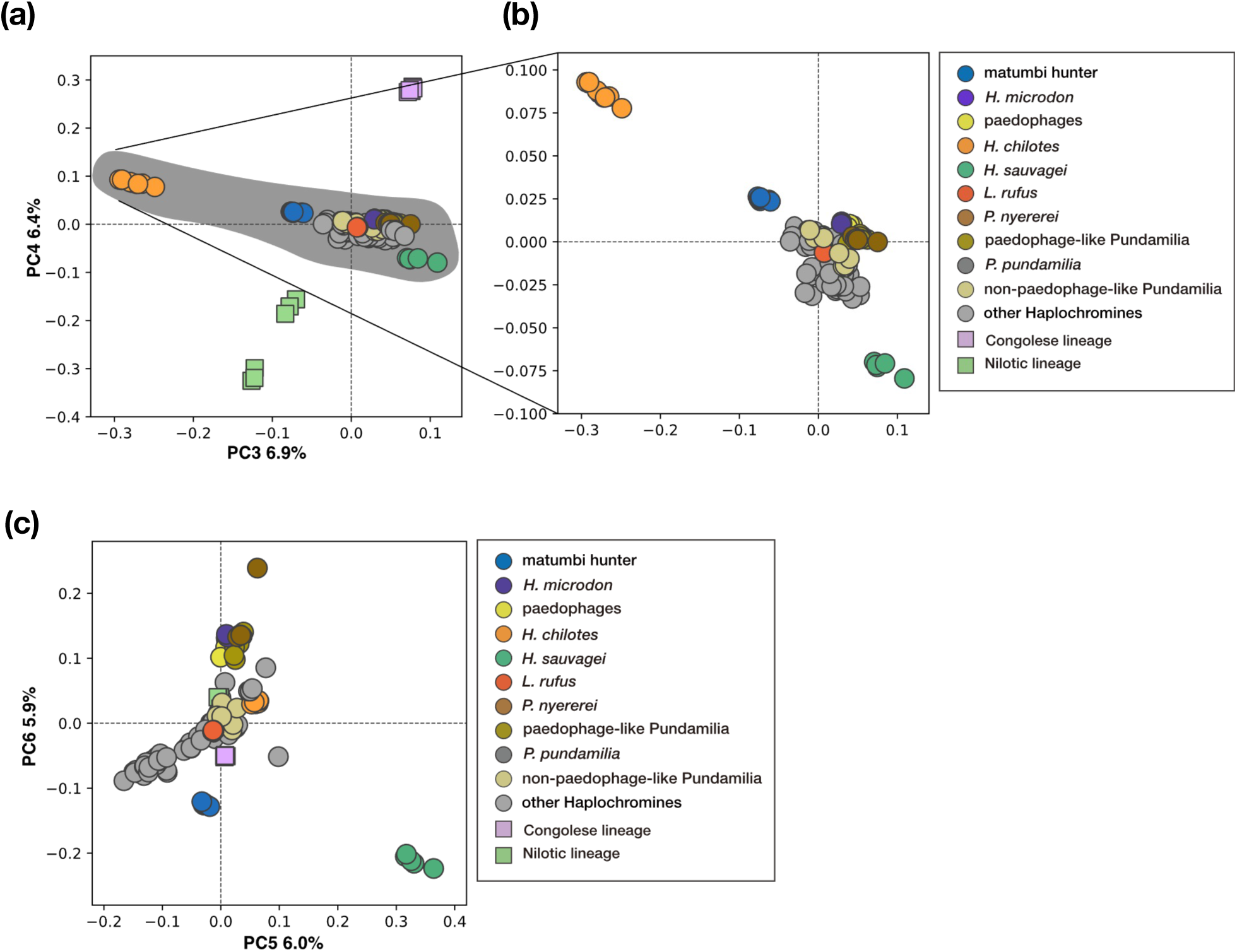
Visualization of population structure in PC3-PC4 (a) and (b) (without outgroup) and PC5-PC6 (c). Samples are labeled with grouping name abbreviations: Nilotic lineage (*Astatotilapia bloyeti*, *A. paludinosa*, *Haplochromis gracilior*, and *Thoracochromis pharyngalis*), Congolese lineage (*A. stappersii*), paedophages (*L. parvidens*, *L. melanopterus*, *L. cryptodon*), paedophage-like Pundamilia (*Pundamilia* sp. “big blue red”, *P*. *igneopinis*, *P*. sp. “nyererei-like”, *P*. sp. “orange”, and *P*. sp. “pundamilia-like”), and non-paedophage-like Pundamilia (*P*. sp. “all red”, *P*. *azurea*, *P*. sp. “blue giant”, *P*. sp. “large red deepwater”, *P*. *macrocephala*, and *P*. sp. “pink anal “). The contribution rate for each PC is written on the axis.

**Figure S3.**
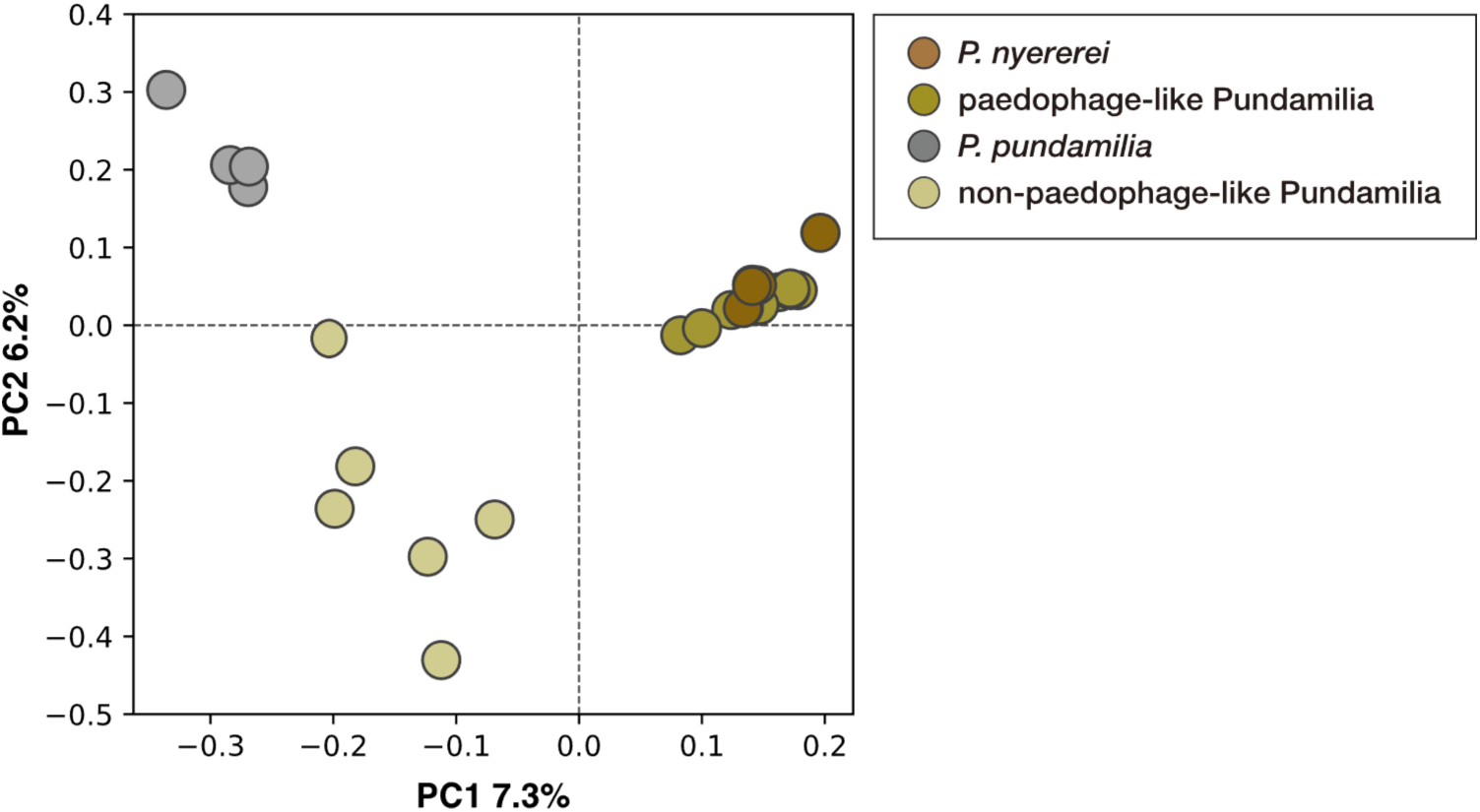
Clustering analysis by PCA for 13 *Pundamilia* species discovered hidden genetic groups within the genus. Species except for P. nyererei and P. pundamilia are labeled with group name: paedophage-like Pundamilia (*Pundamilia* sp. “big blue red”, *P*. *igneopinis*, *P*. sp. “nyererei-like”, *P*. sp. “orange”, and *P*. sp. “pundamilia-like”), and non-paedophage-like Pundamilia (*P*. sp. “all red”, *P*. *azurea*, *P*. sp. “blue giant”, *P*. sp. “large red deepwater”, *P*. *macrocephala*, and *P*. sp. “pink anal “). The contribution rate for each PC is written on the axis.

**Figure S4.**
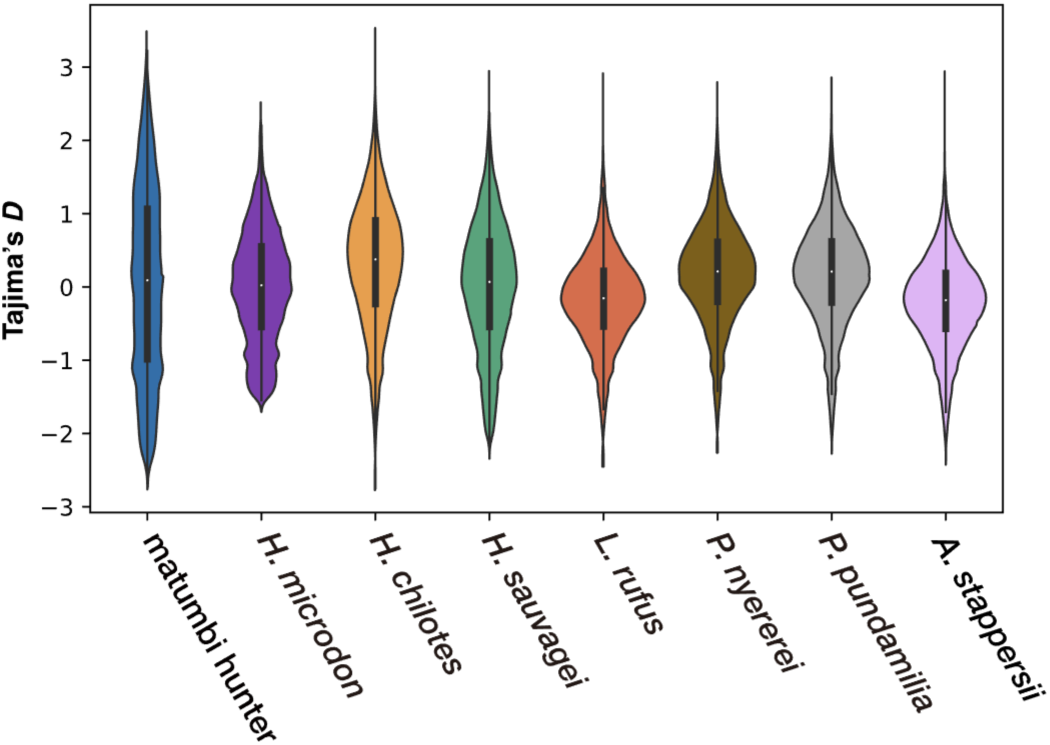
Distribution of Tajima’s *D* calculated by 10,000 bp windows across the whole genome. The range between the first and third quantiles is visualized by a thick line in each violine plot, and the median is depicted by a white circle.

**Figure S5.**
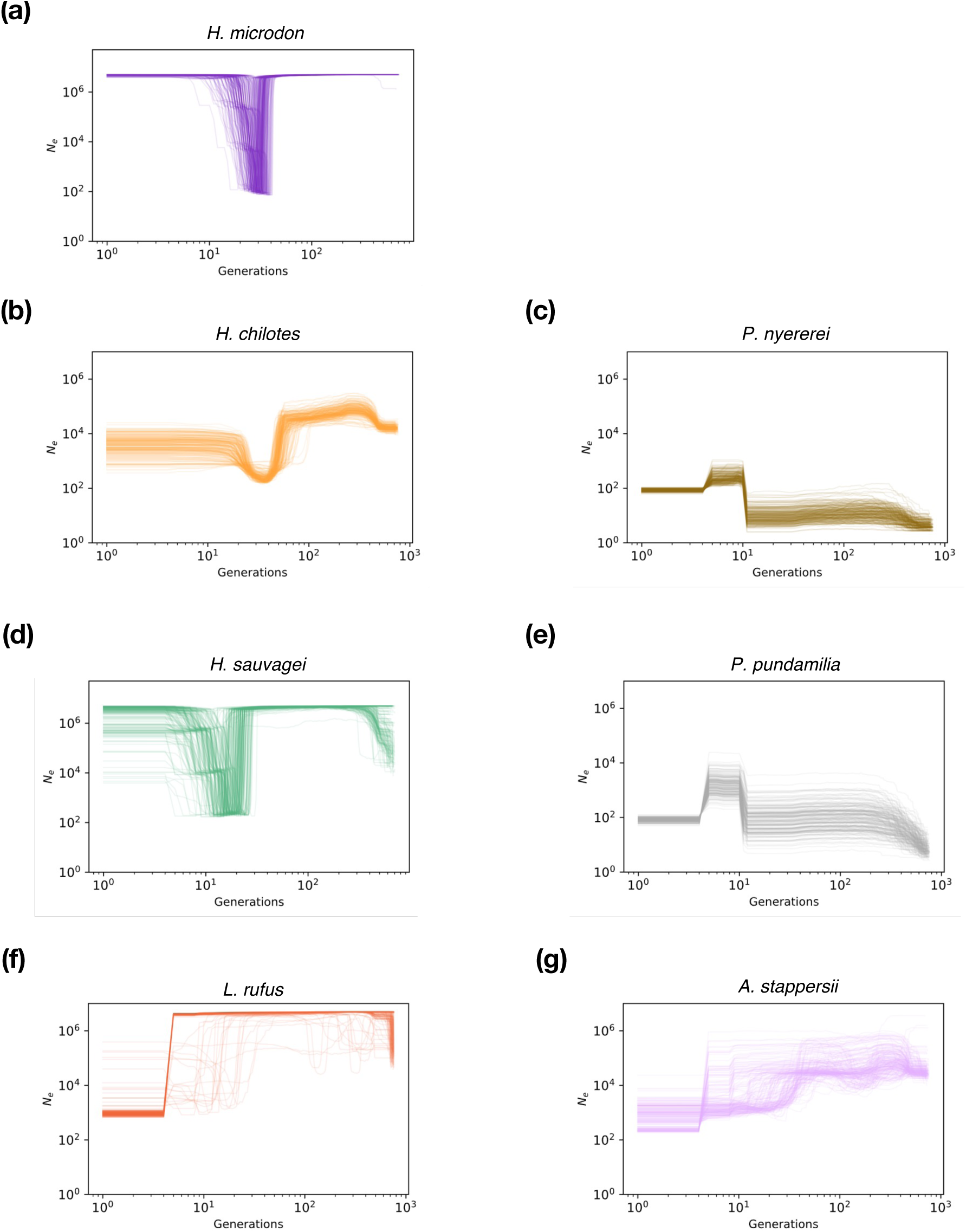
Demographic changes of effective population size (*N*_e_) in the past 700 generations (years) estimated by GONE. The GONE estimate was repeated 200 times and all results were plotted.

**Figure S6.**
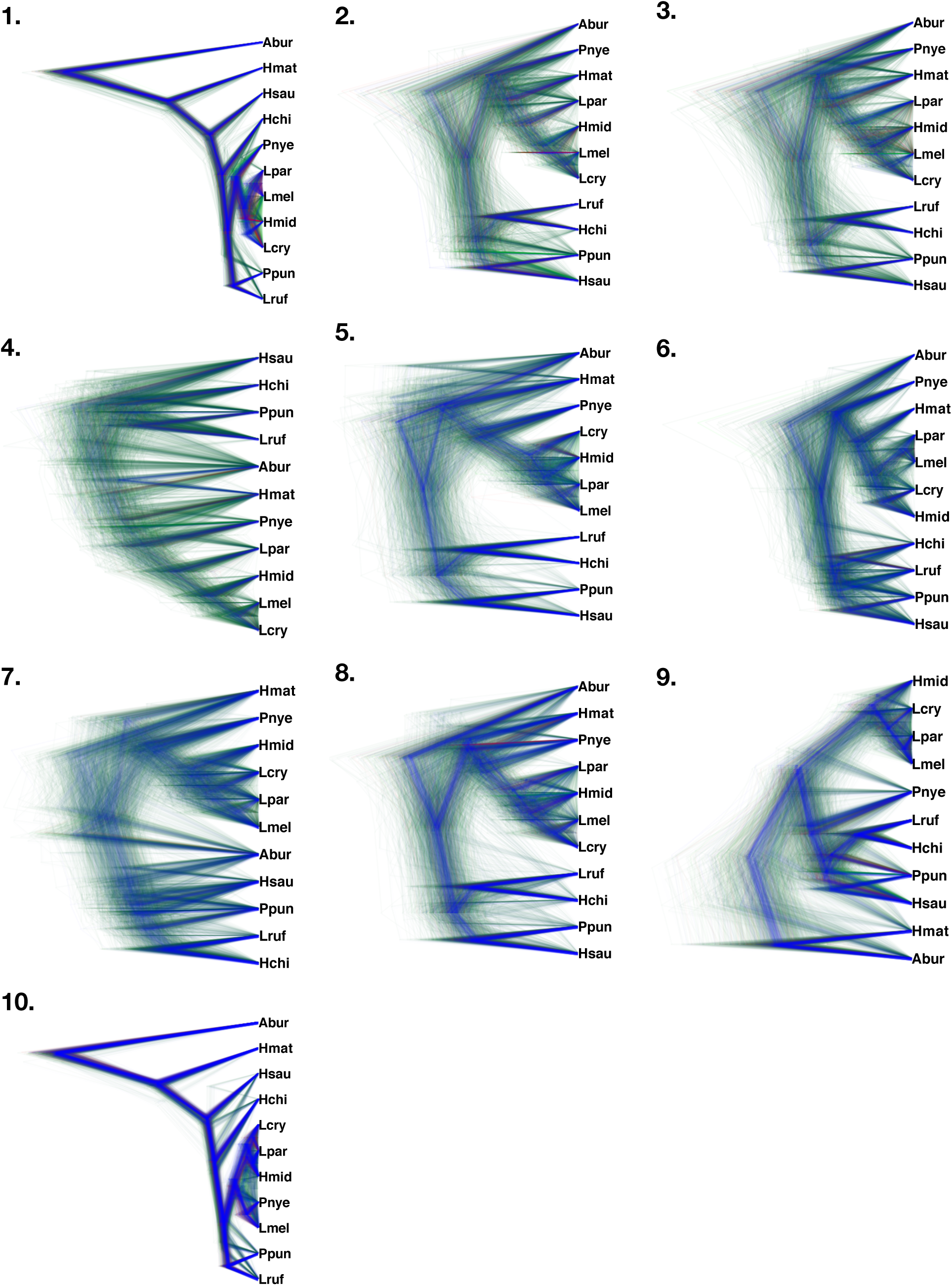
Densitrees constructed by 10 un-concatenated datasets for SNAPP with a consensus topology (ticked blue line). The number in the corner of the trees indicates the dataset number. *Astatotilapia burtoni* (Abur), a riverine lineage, was rooted as an outgroup for all analyses. Samples are labeled with species name abbreviations: Hmat (matumbi hunter), Hmid (*H. microdon*), Lpar (*L. parvidens*), Lmel (*L. melanopterus*), Lcry (*L. cryptodon*), Pnye (*P. nyererei*), Ppun (*P. pundamilia*), Hchi (*H. chilotes*), Hsau (*H. sauvagei*), Lruf (*L. rufus*). The more commonly observed trees are colored in the order of blue, red, and green, while the rest are dark green, respectively.

